# A role for astroglial calcium in mammalian sleep

**DOI:** 10.1101/728931

**Authors:** Ashley M. Ingiosi, Christopher R. Hayworth, Daniel O. Harvey, Kristan G. Singletary, Michael J. Rempe, Jonathan P. Wisor, Marcos G. Frank

## Abstract

Mammalian sleep is characterized by dramatic changes in neuronal activity, and waking neuronal activity is thought to increase sleep need. Changes in other brain cells (glia) across the natural sleep-wake cycle and their role in sleep regulation are comparatively unexplored. We show that sleep is also accompanied by large changes in astroglial activity as measured by intracellular calcium concentrations in unanesthetized mice. These changes in calcium vary across different vigilance states and are most pronounced in distal astroglial processes. We find that reducing intracellular calcium in astrocytes impaired the homeostatic response to sleep deprivation. Thus, astroglial calcium changes dynamically across vigilance states and is a component of the sleep homeostat.

**One Sentence Summary:** Astroglial calcium concentrations vary with sleep and wake, change after sleep deprivation, and mediate sleep need.

## Introduction

Glial astrocytes have been hypothesized to play important roles in mammalian sleep (*1*). They secrete sleep-inducing substances and sleep influences the morphology, gene expression, and proliferation of different glial cells (*1*). Astrocytes are also densely concentrated throughout the brain including regions important for sleep and wake (*2*). They surround synapses, respond non-linearly to neurotransmitters, and via several different mechanisms (*e.g*. metabolism, neurotransmitter uptake, and gliotransmission), modulate neuronal activity (*3–6*). This confers upon these cells the ability to integrate and respond to changes in neurons, providing a feedback mechanism to regulate neuronal activity (*3–5, 7*). There are, however, two critical and unaddressed issues regarding the role of astrocytes in sleep.

A key feature of mammalian sleep is that it is accompanied by widespread and diverse changes in cortical neuronal activity (*8*). In the late 1950’s, this observation led to a revolution in our understanding of sleep. The discovery of rapid eye movement sleep (REMS) dispelled the ‘passive theory’ of sleep according to which the sleeping brain was quiescent (*9*). Sleep has been subsequently discussed and described almost entirely based on neuronal measurements. Astrocytes, however, could be just as dynamic as neurons across the sleep-wake cycle, albeit in different ways. If true, this would trigger a fundamental change in our understanding of sleep by revealing an entirely different level of brain organization that also changes with sleep. This intriguing possibility has not been investigated. Previous sleep studies of astrocytes have relied on *ex vivo* measurements (*10–12*) or did not measure natural patterns of activity in sleep (*13*).

Homeostasis is a second key feature of sleep. In all mammals, sleep intensity (as measured by electroencephalographic (EEG) activity) and/or sleep time increases as a function of prior time awake. Sleep homeostasis has also been historically thought to be primarily a product of neurons (*1*). Recent findings suggest that astrocytes may be part of the mammalian sleep homeostat (*2, 14*), but the cellular mechanisms governing this putative feedback circuit are controversial and largely unexplored (*1*). It is also unknown if essential astroglial signaling pathways [e.g. intracellular calcium (Ca^2+^)] encode sleep need or are essential components of sleep homeostasis. We performed several experiments to investigate these two issues.

## Results

We first determined if astroglial intracellular Ca^2+^ concentrations change across naturally cycling vigilance states. To do this, we expressed the genetically encoded calcium indicator GCaMP6f selectively in frontal cortex astrocytes of C57BL/6J mice (fig. S1). The frontal cortex shows the greatest dynamic range of non-rapid eye movement sleep (NREMS) slow wave (SWA); a canonical index of mammalian sleep need (*2, 15*). We then recorded EEG and electromyographic (EMG) activity while simultaneously imaging changes in astroglial GCaMP6f Ca^2+^ signals in unanesthetized, freely behaving mice using a head-mounted, epifluorescent microscope (Inscopix; Fig. 1A – B, movie S1). We found profound differences in astroglial Ca^2+^ concentrations (Fig. 1C; χ^2^ = 914.39, p < 0.001), frequency of Ca^2+^ events (Fig. 1D; χ^2^ = 295.64, p < 0.001), and event amplitudes (Fig. 1E; χ^2^ = 779.48, p < 0.001). Overall, these astroglial Ca^2+^ measures were greatest in wake and lowest in REMS. An additional striking observation was the transition from REMS to wake, which was accompanied by large changes in Ca^2+^ concentrations (Fig. 1F, movie S1). Sleep is also accompanied by changes in the synchrony of neuronal networks (*8*). It was therefore of interest to explore synchronous activity in astrocytes during different vigilance states. As shown in fig. S2, astrocytes were highly synchronized in all states, but this synchrony was greatest during wake and lowest in REMS. Collectively, these data show that at the level of single cells and networks, astrocytes (like neurons) display dramatic changes in activity across the sleep-wake cycle.

**Fig. 1.**
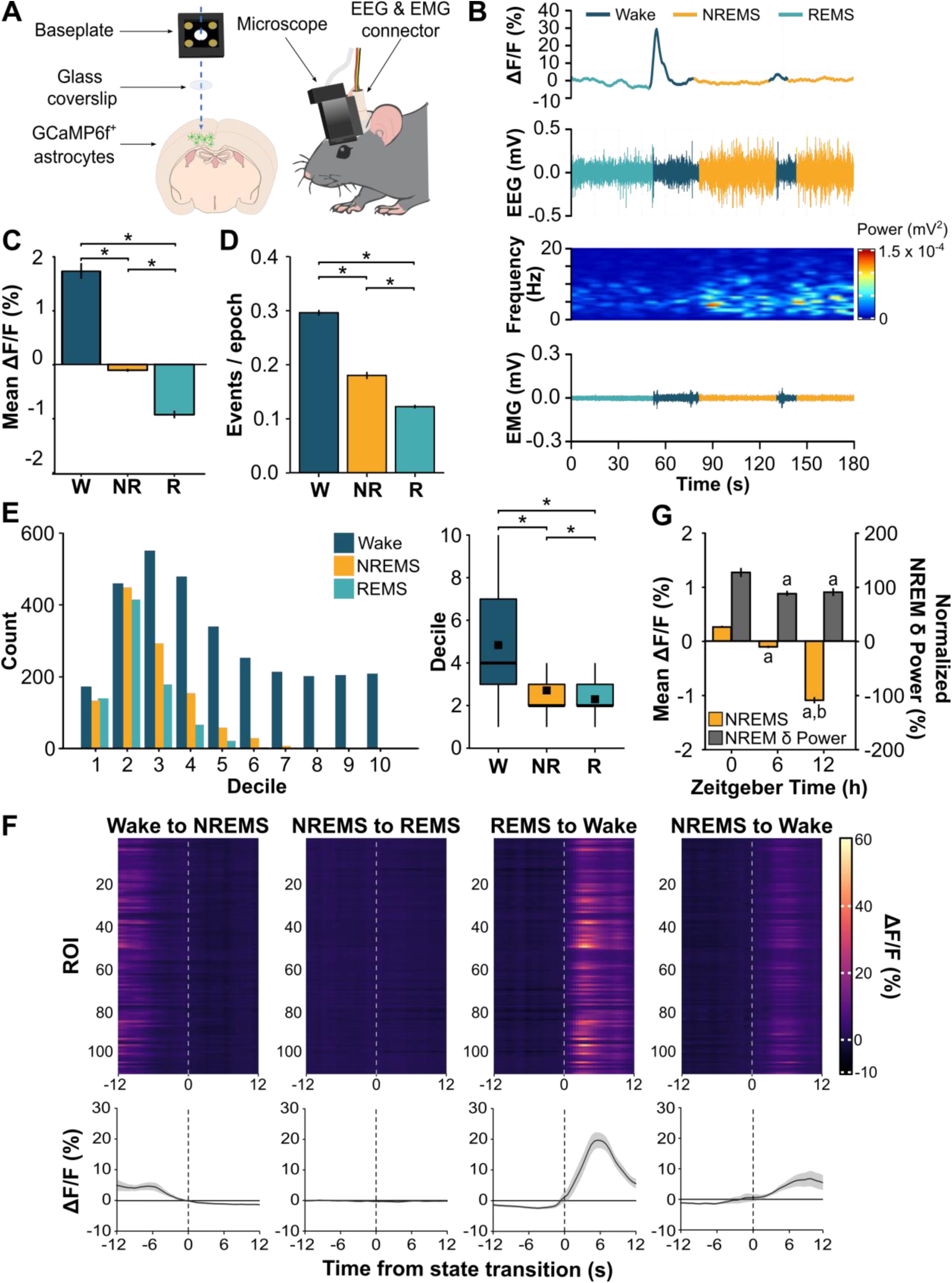
Astroglial Ca^2+^ dynamics change with vigilance state and time of day. (**A**) Cartoon showing dual EEG & EMG recording and astroglial Ca^2+^ imaging in the frontal cortex through a cranial window. (**B**) Representative Ca^2+^-induced fluorescent trace, EEG, EEG power spectrogram, and EMG in each vigilance state. (**C**) Mean (± s.e.m.) ΔF/F during wake (W), NREMS (NR), and REMS (R) at ZT6. (**D**) Number of astroglial Ca^2+^ events per 4-s vigilance state epoch expressed as means ± s.e.m. at ZT6. (**E**) Distribution of Ca^2+^ event amplitudes organized by smallest (low deciles) to largest (high deciles) values during wake, NREMS, and REMS and boxplot comparisons at ZT6. Black squares in boxplots denote means. (**F**) Heatmaps show fluorescence traces from 110 regions of interest (ROI) from a representative mouse during the 12 s before and after transitions between vigilance states. Line plots are mean ΔF/F (± s.e.m.) during state transitions under baseline conditions for n = 6 mice. Vertical dashed lines indicate time of state transition. (**G**) Mean ΔF/F during NREMS and baseline NREMS EEG delta (δ) power (0.5 – 4 Hz) at ZT0, 6, and 12. NREM δ power is expressed as a percentage of the last 4 h of the light period. Values are means ± s.e.m. ^a^, different from ZT0; ^b^, different from ZT6 (linear mixed-effects model). Group means are from n = 586 ROIs from 6 mice. *, vigilance state differences (Friedman test). p < 0.05.

We next explored whether astroglial Ca^2+^ concentrations change across the rest phase. This would provide insight into whether astrocytes encode sleep need because sleep need (as measured by NREMS SWA) is high at the beginning of the rest phase (Zeitgeber time [ZT] 0) and discharged by the end (ZT12) (*16, 17*). We found that mean NREMS GCaMP6f fluorescence was maximal at ZT0 and reached a nadir at ZT12 (F(2, 1566.18) = 85.93, p < 0.001), in parallel with changes in NREMS SWA (Fig. 1G; F(2, 387.07) = 9.13, p < 0.001). Changes in wake and REMS, however, were more variable over time (fig. S3). These results suggest that astrocytes change their activity in parallel with circadian variations in sleep need and this is most consistently found in NREMS.

We then determined if changes in Ca^2+^ concentration were uniform throughout the astrocyte or compartmentalized in the soma vs. the distal processes. This was of interest because Ca^2+^ dynamics in astroglial processes can occur independently from the soma and may reflect different types of signal processing (*18–21*). Because the Inscopix signal includes both somatic and distal process activity, we used two-photon microscopy to more precisely examine how Ca^2+^ dynamics change in these regions across vigilance states in unanesthetized mice (Fig. 2A – B, movie S2). Distal processes showed greater frequency of single Ca^2+^ events compared to somata in each vigilance state (Fig. 2C; wake: U = 41,221.00, p < 0.001; NREMS: U = 45,6161.50, p < 0.001; REMS: U = 45,023.00, p < 0.001). Furthermore, fluorescent intensity was greater in astroglial processes compared to somata during wake (Fig. 2D; U = 99,967.50, p < 0.001) and NREMS (U = 87,228, p < 0.001). REMS exhibited a similar trend (U = 63,555.00, p = 0.102). However, measuring the difference between somatic and distal process Ca^2+^ concentrations revealed that the bias towards distal processes was largest in wake and REMS (Fig. 2E; χ^2^ = 267.57, p < 0.001). Overall, these findings indicate that astroglial activity is not uniform during sleep but biased to distal processes. This suggests that sleep is accompanied by astroglial sampling of local synaptic activity and/or changes in microdomain metabolism (*19–21*).

**Fig. 2.**
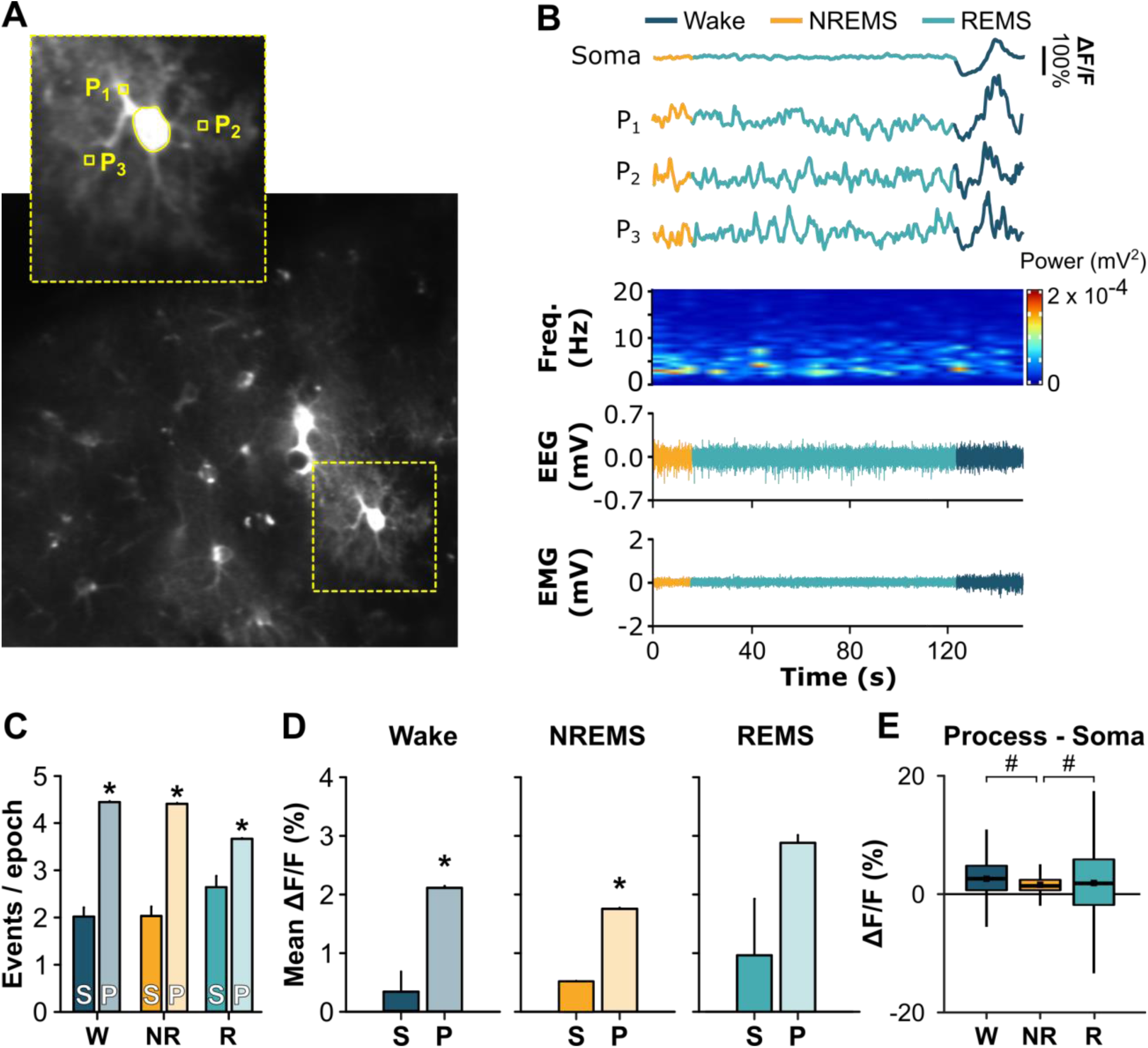
Astroglial Ca^2+^ dynamics differ between somata and processes. (**A**) Sum z-stack projection image of GCaMP6f^+^ astrocytes from two-photon microscopy. Inset shows regions of interest for soma and distal processes (P_1_ – P_3_) for a single astrocyte depicted in **B**. (**B**) Representative Ca^2+^ traces from an astrocyte soma and 3 of its respective processes (P_1_ – P_3_) with aligned EEG power spectrograms, EEG, and EMG. (**C**) Number of Ca^2+^ events per 4-s vigilance state epoch in somata (S) and processes (P) expressed as means ± s.e.m. under baseline conditions for wake (W), NREMS (NR), and REMS (R) during the rest phase (Mann-Whitney U). (**D**) Mean ΔF/F (± s.e.m.) for wake, NREMS, and REMS for astroglial somata (S) and processes (P) under baseline conditions during the rest phase (Mann-Whitney U). (**E**) Difference scores for ΔF/F between process ROIs and their respective soma for W, NR, and R during the rest phase (Friedman). Black squares in boxplots denote means. Group means are from n = 60 somata and n = 6074 processes from 4 mice. *, different from somata. #, difference between vigilance states. p < 0.05.

We then more directly tested whether changes in astroglial Ca^2+^ encode changes in sleep need. Although this was suggested by our time of day measures (Fig. 1G), this is traditionally explored using sleep deprivation (SD). To do this, we sleep deprived freely behaving mice for 6 h (ZT0 – 6) and imaged using the Inscopix system. SD increased NREMS SWA (175.34 ± 5.05% vs. baseline) as expected but also induced state-dependent changes in astroglial Ca^2+^. At ZT6 (when sleep need is maximally discharging immediately after SD (*16, 17*)) there was an overall increase in mean Ca^2+^ concentrations in all states (Fig. 3A; wake: Z = −10.47, p < 0.001; NREMS: Z = −3.47, p = 0.001; REMS: Z = −9.74, p < 0.001). In contrast, the frequency of Ca^2+^ events increased in wake but decreased in both sleep states (Fig. 3B; wake: Z = −9.21, p < 0.001; NREMS: Z = −19.77, p < 0.001; REMS: Z = −18.95, p < 0.001). We also determined if SD changed the network synchrony of cortical astrocytes. Interestingly, and in contrast to cortical neurons (*22*), SD reduced synchronous activity in NREMS, but increased it in REMS (fig. S2B). These results thus confirmed that astroglial Ca^2+^ changes in parallel with sleep need, consistent with the idea that this signal encodes sleep need. The data further showed that these changes were distinct from those reported in electrophysiological studies of neurons (*22*), suggesting that the former were not a passive response to surrounding neuronal activity.

**Fig. 3.**
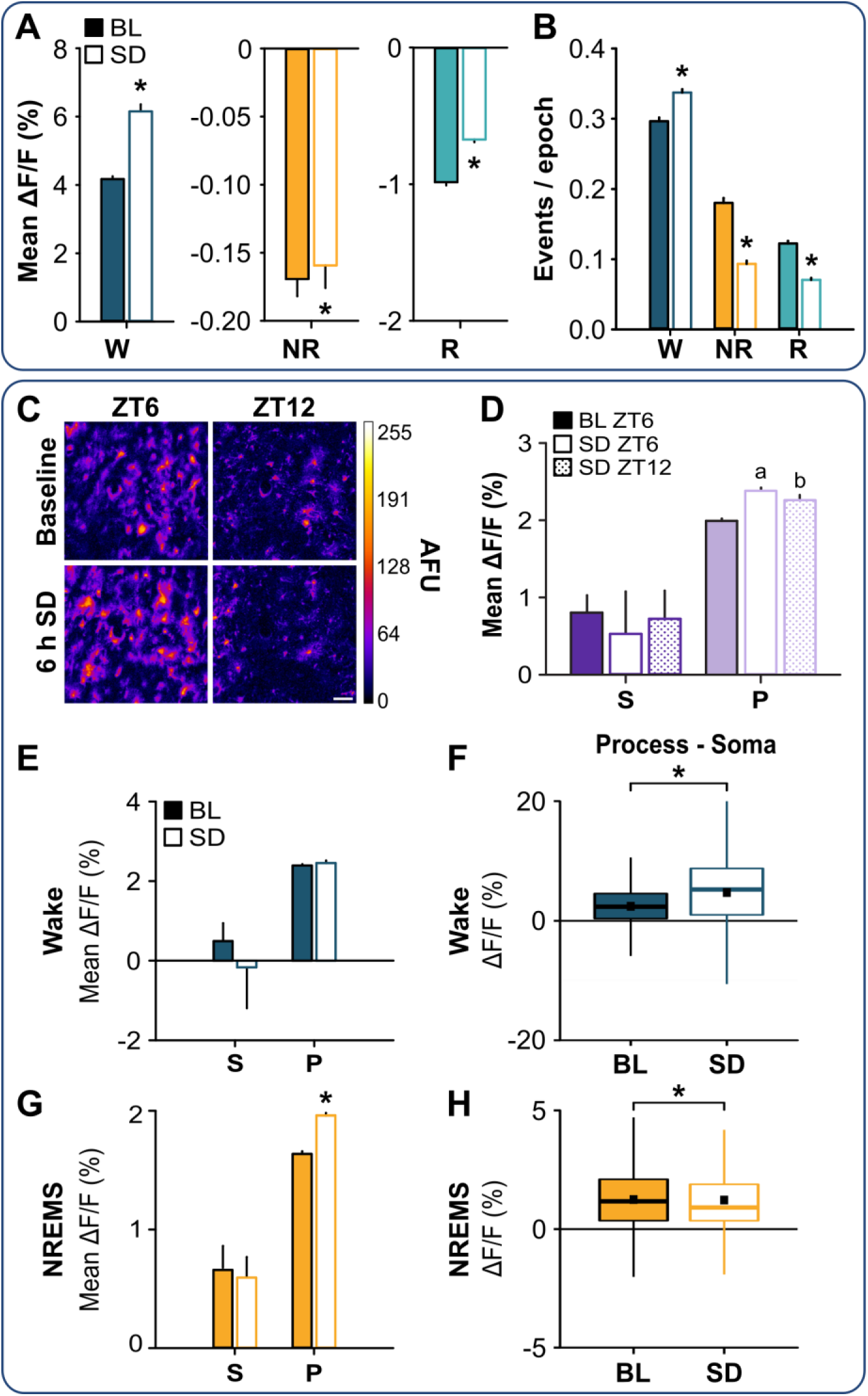
Sleep deprivation alters astroglial Ca^2+^ dynamics. Upper box (**A** – **B**) shows data from head-mounted epifluorescent microscope (Inscopix). Group means are from n = 586 somata from 6 mice. Lower box (**C** – **H**) shows data from two-photon imaging. Group means are from n = 60 somata and n = 6074 processes from 4 mice. (**A**) Mean ΔF/F values at ZT6 during wake (W), NREMS (NR), and REMS (R) under baseline (BL; closed bars) and sleep deprived (SD; open bars) conditions (Wilcoxon signed rank). (**B**) Mean number of astroglial Ca^2+^ events per 4-s vigilance state epoch at ZT6 during W, NR, and R under BL and SD conditions (Wilcoxon signed rank). (**C**) Maximum projection images of GCaMP6f^+^ astrocytes at ZT6 (left) & ZT12 (right) for BL (top) and SD (bottom). Scale bar = 50 μm. Color bar shows arbitrary fluorescence units (AFU). (**D**) Mean ΔF/F values from all vigilance states for somata (S) and processes (P) during BL ZT6, SD ZT6, and SD ZT12. ^a^, difference from BL ZT6 processes; ^b^, difference from SD ZT6 processes (Friedman). (**E** & **G**) Mean ΔF/F values during (**E**) wake and (**G**) NREM during ZT6 under BL and SD conditions (Wilcoxon signed rank). (**F** & **H**) Distributions of differences in ΔF/F values between processes and their soma during ZT6 under BL and SD conditions for (**F**) wake and (**H**) NREMS (Wilcoxon signed rank). Black squares in boxplots denote means. Values are means ± s.e.m. for A, B, D, E and G. *, different from BL. p < 0.05.

We used two-photon microscopy to more finely explore the spatial and temporal aspects of this compensatory Ca^2+^ response to SD (Fig. 3C). Averaged across all states, changes in Ca^2+^ concentrations were greatest in the processes following sleep deprivation (Fig. 3D; processes: χ^2^ = 106.28, p < 0.001). Mean Ca^2+^ concentrations at ZT6 in the processes was elevated after SD relative to baseline then declined at ZT12; this paralleled the discharge of NREM SWA from ZT6 (187.55 ± 20.56%) to ZT12 (106.00 ± 10.51%). We then examined within-state changes. Wake was accompanied by a trend towards lower somatic Ca^2+^ concentrations (Fig. 3E, fig. S4) which contributed to a significant shift in Ca^2+^ concentrations towards the processes (Fig. 3F; Z = −29.37, p < 0.001). During NREMS, Ca^2+^ concentrations in processes were elevated after SD at ZT6 (Fig. 3G; Z = −16.05, p < 0.001; Fig. 3H; Z = −6.53, p < 0.001) and reached baseline levels at ZT12 (fig. S4A). Ca^2+^ dynamics during REMS were more variable (fig. S4, fig. S5). Thus, more fine-grained analyses revealed that SD-induced changes in astroglial Ca^2+^ were most consistently found in NREMS and primarily reflected increases at distal processes.

**Fig. 4.**
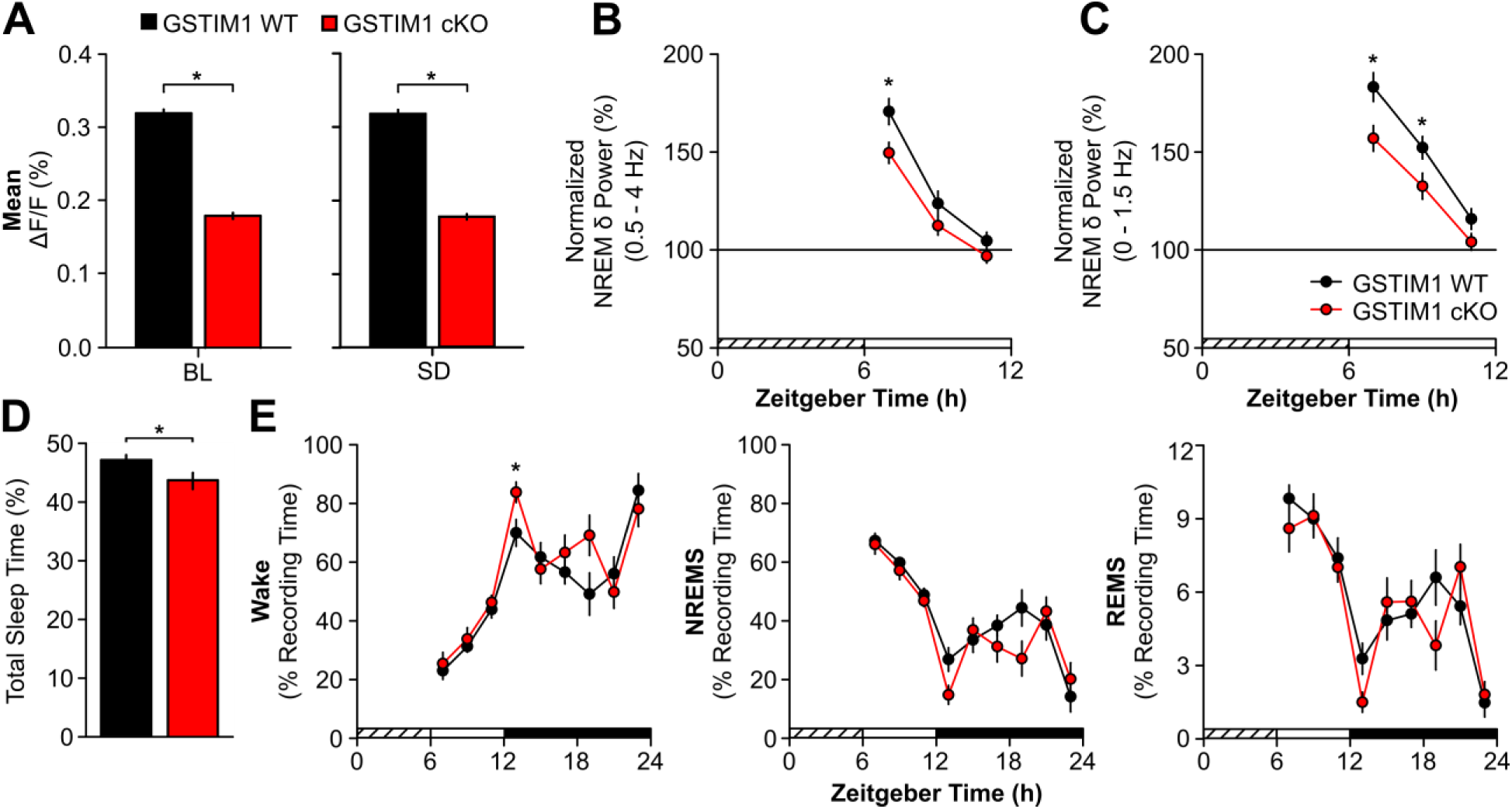
Reductions in astroglial Ca^2+^ attenuate homeostatic response to sleep deprivation. **A**) Mean ΔF/F values for GSTIM1 WT (black) and GSTIM1 cKO (red) at ZT6 under baseline (BL) and sleep deprived (SD) conditions (Mann-Whitney U). (**B**) Normalized NREM δ power (0.5 – 4 Hz) and (**C**) normalized low NREM δ power (0 – 1.5 Hz) during the first 6 h of recovery post-SD expressed in 2 h bins (repeated measures ANOVA). (**D**) Total time spent asleep (sum of NREMS and REMS) during the 18 h recovery phase post-SD expressed as percentage of total recording time (unpaired t-test). (**E**) Time spent in wake (left), NREMS (middle), and REMS (right) during the 18 h recovery phase post-SD expressed as a percentage of recording time across 2 h bins (repeated measures ANOVA). Values are means ± s.e.m. *, genotypic differences. p < 0.05.

The time of day and SD experiments strongly suggested that astroglial Ca^2+^ encodes sleep need. To more directly determine if astroglial Ca^2+^ was required for sleep homeostasis, we inducibly deleted stromal interaction molecule 1 (STIM1) in astrocytes. STIM1-mediated store-operated Ca^2+^ entry (SOCE) is an essential mechanism by which the intracellular Ca^2+^ concentration is elevated (*23*). Inhibiting STIM1 impairs SOCE in astrocytes (*23*), indicating that STIM1 is an integral component of astroglial Ca^2+^ regulation. We first determined that astroglial Ca^2+^ concentrations were lower in mice lacking STIM1 in astrocytes (GSTIM1 conditional knockout [cKO]) compared to GSTIM1 wild type (WT) mice (Fig. 4A; BL: U = 12,236.00, p < 0.001; SD: U = 11,866.00, p < 0.001; see also fig. S6). Next, we tested if reduced astroglial Ca^2+^ concentrations affected sleep and wake behavior under baseline conditions and after 6 h SD. Interestingly, GSTIM1 WT and cKO mice were largely similar for time spent in each vigilance state, bout frequency, bout duration, and EEG spectral power (fig. S7). After SD, however, the normal homeostatic response to SD was blunted in GSTIM1 cKO mice as measured by NREM SWA (0.5 – 4.0 Hz; Fig. 4B; genotype x time effect, F(2,26) = 4.00, p = 0.031and Fig. S8), NREM SWA in the lower frequency bands (0.0 – 1.5 Hz; Fig. 4C; genotype effect, F(1,13) = 7.00, p = 0.020), and sleep time (Fig. 4D; t(15) = 2.23, p = 0.041 and Fig. 4E; genotype effect for wake, F(1,15) = 4.98, p = 0.041). These findings demonstrate that astroglial STIM1-mediated Ca^2+^ is a key mediator of sleep homeostasis.

## Discussion

We investigated the role of astroglial Ca^2+^ in mammalian sleep. We find that this essential mediator of astroglial function changes dramatically across sleep and wake and after SD. Reducing the astroglial Ca^2+^ signal *in vivo* also reduces the homeostatic response to SD. These findings demonstrate that sleep is not only accompanied by widespread activity changes in neurons, but also by changes in glial cells. They further suggest that astroglial Ca^2+^ signaling is part of the mammalian sleep homeostat.

Our findings address several important issues in the field of sleep and glial biology. First, they suggest that during sleep there is a separate level of brain organization (glia) that changes as dynamically as neurons. If true, this would fundamentally change our scientific understanding of sleep, because this has been shaped almost entirely by neuron-based studies. One possible explanation for our findings is that changes in astroglial Ca^2+^ are passively driven by changes in surrounding, local neuronal activity. For example, in all vigilance states, most of the astroglial Ca^2+^ signal was in distal processes that surround and respond to changes in local synaptic activity and metabolism (*18–21*). On the other hand, astroglial Ca^2+^ did not appear to merely mirror patterns of neuronal activity (reported with electrophysiology (*8, 22*); but see (*24*)) characteristic of wake and sleep. For example, wake and REMS display similar levels and patterns of cortical neuron firing (*8, 22*) but this was not true for astroglial Ca^2+^. In addition, while NREMS is typified by highly synchronous neuronal activity, which is increased after SD (*8, 22*), synchrony in astrocytes in NREMS was lower than in wake and reduced even further after SD. This raises the possibility that the regulation and function of sleep may more directly involve glial cells.

Second, our findings support the idea that astrocytes play a role in sleep homeostasis (*2*). SD in mammals produces homeostatic and compensatory changes in neurons (*22, 25*). We now show that an entirely different class of brain cells also exhibits compensatory changes in response to SD. Across all vigilance states, Ca^2+^ changes in NREMS were most aligned with the accumulation and discharge of sleep need as measured by NREMS SWA. This is notable because the need for sleep is best modeled by changes in NREMS (*16, 17*). Conditional reduction of this signal also selectively reduced classic measures of sleep homeostasis (NREMS SWA and sleep time). Similar results were reported following inducible inhibition of astrocytic gliotransmission, particularly in the lower SWA bands that are most sensitive to sleep loss (*2*). This suggests that changes in astroglial intracellular Ca^2+^ encode sleep need and trigger feedback onto neurons perhaps via gliotransmission of ATP (*19*), which is hydrolyzed to the sleep-inducing molecule adenosine (*2*).

Lastly, our findings address an ongoing controversy in the field of glial biology. Astroglial Ca^2+^ dynamically changes in response to neurotransmitters, metabolic signals and mechanical stimulation. Studies *in vitro* have also shown that increases in Ca^2+^ trigger gliotransmission (*26, 27*). Yet, previous manipulations of intracellular astroglial Ca^2+^ *in vivo* have failed to reveal significant behavioral phenotypes (*28*). We used a novel approach to this problem (GSTIM1 cKO mice) that targeted a rate-limiting step in ER replenishment of astroglial intracellular Ca^2+^. We find that this manipulation selectively alters mammalian sleep homeostasis, thus demonstrating a functional role for astroglial Ca^2+^ in a complex, regulated brain state *in vivo*.

## Acknowledgements

We thank Taylor Wintler and Hannah Schoch for assisting with SD. We also thank Will Clegern for engineering and fabricating custom hardware for imaging and Eric Marr for assistance in analysis. We thank Mark Wu for discussions of unpublished results.

## Author contributions

Conceptualization, A.M.I. and M.G.F.; Experimental design; A.M.I. and M.G.F.; Investigation, A.M.I. and K.G.S.; Analysis, A.M.I., C.R.H., D.O.H., J.P.W., M.J.R., and M.G.F., Writing – original draft, A.M.I., C.R.H., D.O.H, and M.G.F. Writing – review & editing, A.M.I., C.R.H., D.O.H., K.G.S., M.J.R., J.P.W., and M.G.F.

## Data and materials availability

Data are password protected and backed up on a secure server. Data are available upon request.

## Materials and Methods

### Animals

C57Bl/6J (#000664), B6.Cg-Tg(GFAP-cre/ERT2)505Fmv/J (GFAP-Cre/ERT2; #12849), and B6(Cg)-Stim1^tm1Rao^/J (STIM1^fl/fl^; # 023350) mice were obtained from The Jackson Laboratory (Bar Harbor, ME). Breeding pairs of hemizygous GFAP-Cre/ERT2^Tg+/-^ and homozygous STIM1^fl/fl^ were established to obtain GFAP-Cre/ERT2^-/-^;STIM1^fl/fl^ wild type mice (GSTIM1 WT) and GFAP-Cre/ERT2^Tg+/-^;STIM1^fl/fl^ conditional knockout (GSTIM1 cKO) mutant mice. Mice were housed in standard cages on 24 ± 1°C on a 12:12 h light:dark cycle with food and water *ad libitum*. All experimental procedures were approved by the Institutional Animal Care and Use Committee of Washington State University and conducted in accordance with National Research Council guidelines and regulations for experiments in live animals.

### Surgical Procedures

#### Ca^2+^ imaging with EEG & EMG recording

Adult male mice (8 – 14-weeks-old) were anesthetized using isoflurane and placed in a stereotaxic frame. A 3-mm craniotomy was made over the frontal cortex leaving the dura intact. AAV2/5 *GfaABC_1_D*-GCaMP6f (3.31 x 10^13^ GC/ml; Penn Vector Core, Philadelphia, PA) was injected at two adjacent sites (AP: 2.0 - 2.5 mm, ML: - 1.25 – 1.75 mm, DV: −0.18 – 0.20 mm) in the frontal cortex. 1.5 µl of vector was injected at each site at 200 nl/min. The needle remained in place for 10 min after the injection was complete. After vector delivery, a 3-mm glass coverslip was fixed over the craniotomy with cyanoacrylate adhesive. For electrode implantation, four 0.5-mm diameter holes were drilled in the skull unilaterally over frontal, somatosensory, and occipital cortices in the hemisphere contralateral to the AAV2/5 *GfaABC_1_D*-GCaMP6f injection site. Sterilized stainless steel EEG electrodes were inserted into these holes just under the skull, and 2 additional EMG electrodes were placed into the nuchal muscles. Cranial electrodes were secured with dental acrylic. For two-photon imaging, a custom-milled head-restraint bar was then fixed to the skull with dental acrylic mixed with black carbon powder. Mice were imaged 2 – 4 weeks later to allow time for the fluorescent indicator to be expressed (*18, 29*).

For imaging with the head-mounted microscope, mice were fitted with a baseplate 1.5 – 2.5 weeks after surgery under isoflurane anesthesia. The baseplate used to house the microscope was secured with dental acrylic mixed with black carbon powder.

#### EEG & EMG implantation for sleep phenotyping

Adult male mice (10 – 14-weeks-old) were stereotaxically implanted with EEG & EMG electrodes under isoflurane anesthesia according to previously published methods (*30, 31*). Briefly, four stainless steel screw electrodes (BC-002MPU188, Bellcan International Corp, Hialeah, FL) were placed contralaterally over frontal (2) and parietal (2) cortices, and 2 EMG electrodes were inserted in the nuchal muscles. Mice were allowed 5 days of recovery from surgery prior to habituation to the recording environment.

### Immunofluorescence

C57Bl/6J mice expressing GCaMP6f in frontal cortex were transcardially perfused with 10% buffered formalin. Brains were extracted and placed in 30% sucrose in phosphate buffered saline (PBS) for 24 – 48 h. Brains were frozen in −50°C 2 -methylbutane and stored at −80°C until processing. Brains were sectioned coronally at 25 µm on a cryostat. Native green fluorescent protein (GFP) signal from GCaMP6f expression was assessed, and sections were also counterstained for astrocytes and neurons using immunofluorescent techniques. Sections were washed 3 x 15 min in PBS and then incubated overnight at room temperature with primary antibodies against astrocytes (1:1000; mouse anti-GFAP, #3670, Cell Signaling Technology, Danvers, MA) or neurons (1:100; mouse anti-NeuN, #MAB377, Millipore, Burlington, MA) in 1x PBS with 0.3% Triton-X (PBSt). The next day, sections were washed 3 x 10 min in PBS and then incubated for 1.5 h at room temperature with biotinylated horse anti-mouse (1:1000; #BA-2000, Vector Laboratories, Burlingame, CA) secondary antibody in PBSt. Sections were again washed for 3 x 10 min in PBS and then incubated for 2 h at room temperature in Texas Red streptavidin (1:500; #SA-1200, Vector Laboratories) in PBSt. After 3 x 10 min washes in PBS, sections were mounted on slides for imaging.

### Experimental Design

#### Ca^2+^ imaging with EEG & EMG recording

Adult male C57Bl/6J mice underwent a counterbalanced design of 24 h undisturbed baseline sleep and 6 h SD via gentle handling followed by 18 h recovery sleep to account for potential order effects of imaging over multiple days. For epifluorescent microscopy (n = 6), EEG & EMG were continuously recorded and imaging occurred at zeitgeber times (ZT) of high (ZT0) and low (ZT6 and ZT12) sleep need. At ZT0 and ZT12, images were acquired during a 2-min imaging session every 20 min. At ZT6, our primary analysis timepoint, images were acquired during 2 – 7 min imaging sessions at varying intervals to capture all vigilance states and state transitions. For two-photon microscopy (n = 4), imaging and EEG & EMG recordings occurred at ZT6 and ZT12 during 3-min imaging sessions at varying intervals to capture all vigilance states. During imaging experiments, vigilance states were determined by visual inspection of the EEG & EMG signals in real time. For epifluorescent imaging in GSTIM1 mice (n = 3 GSTIM1 WT; n = 4 GSTIM1 cKO), EEG & EMG recordings occurred continuously during a 24 h undisturbed baseline day followed by 6 h SD via gentle handling and 18 h recovery sleep. Ca^2+^ imaging occurred at ZT0, 6, and 12 as described for C57Bl/6J mice.

#### GSTIM1 sleep phenotyping

Male and female GSTIM1 WT (n = 8; females = 1) and GSTIM1 cKO (n = 9; females = 4) adult mice (8 – 12-weeks-old) were injected with tamoxifen (180 mg/kg; #T5648, Sigma-Aldrich, St. Louis, MO) intraperitoneally once per day for 5 consecutive days alternating sides (*14*). Tamoxifen was made up at 30 mg/ml in 90% sunflower oil (#S5007, Sigma-Aldrich) with 10% ethanol and sterile filtered through a 0.22 µm filter (*14*). All mice received 1 ml lactated Ringer’s solution with 5% dextrose subcutaneously daily until mice started regaining body weight. Ten days after the last tamoxifen injection, mice underwent surgery for EEG & EMG electrode implantation. Mice were allowed 5 days of recovery from surgery. Animals were then connected to a lightweight, flexible tether and allowed at least 3 days to habituate to the tether and recording environment. After habituation, mice underwent 24 h undisturbed baseline EEG & EMG recording starting at light onset (ZT0). The next day, mice were sleep deprived for 6 h via gentle handling beginning at light onset according to previously published methods (*2, 32*). Mice were then allowed 18 h undisturbed recovery sleep. There were no differences between male and female mice for our sleep measures of interest with two exceptions. Female GSTIM cKO mice had fewer NREMS bouts during the BL light period (male: 9.37 ± 0.36, female: 7.79 ± 0.40; repeated measures ANOVA, effect of sex: F (1, 7) = 9.87, p = 0.016), and there was an effect of sex on BL REMS bout duration for GSTIM cKO mice (repeated measures ANOVA: F (1, 7) = 9.86, p = 0.016) but no significant pairwise differences.

### Chronic EEG & EMG recording with epifluorescent Ca^2+^ imaging in freely behaving mice

#### Data acquisition

Mice were connected to a counterbalanced, light-weight EEG & EMG recording tether coupled to a nano connector (#A79108-001, Omnetics, Minneapolis, MN) and a miniaturized, head-mountable epifluorescent microscope (nVista 2.0; Inscopix, Palo Alto, CA) coupled to a baseplate affixed to the skull. Animals were allowed at least 3 days to habituate to the recording environment. EEG & EMG data were acquired through Pinnacle Technologies Data Conditioning and Acquisition System (#8401 HS) and Sirenia Acquisition software (v1.7.7, Pinnacle Technologies, Lawrence, KS). EEG data were digitized at 250 Hz, high pass filtered at 0.5 Hz, and low pass filtered at 40 Hz. EMG data were digitized at 2 kHz, high pass filtered at 10 Hz, and low pass filtered at 100 Hz. The system was set up to detect rising transistor-transistor logic (TTL) pulses generated by the imaging system, which was used to synchronize imaging frames and electrophysiological data.

Ca^2+^ imaging data was acquired through the miniaturized, head-mountable epifluorescent Inscopix microscope and Inscopix nVista HD software (v2.0.4). Imaging frames were captured at 10.1 frames per second with an exposure time of 49.664 ms at a gain of 2.0. LED power ranged from 50 – 80% to adjust the upper tail of the histogram to be as close to a pixel value of 1500 to ensure good signal-to-noise ratio (*33*). The LED power was set at the beginning of the experiment for each mouse and did not change for the remainder of the experiment. During image capture, TTL pulses were sent to the Sirenia Acquisition software at 50% duty and recorded as annotations in the EEG & EMG data file. The microscope remained in place in the baseplate for the entirety of the EEG & EMG recording period.

#### Data processing and analysis

Imaging data were processed using the Data Processing Software (Inscopix). For each animal, movies acquired during baseline and during SD and recovery were concatenated into a timeseries. This timeseries was spatially downsampled by a factor of 2 from 1440 x 1080 pixels to 720 x 540 pixels and temporally downsampled by a factor of 5 to 2 Hz to reduce the data footprint (*33*). Defective pixels were also rectified (*33*). Small lateral displacements were then corrected for by registering all frames to a single reference frame free of movement using the motion correction algorithm in the Data Processing Software. To identify individual cells, images were converted to ΔF/F images by expressing each frame as the relative change from the mean image obtained from the entire movie (ΔF/F_mean_ = F - F_mean_/F_mean_) (*33, 34*). A spatial bandpass filter was also applied using default settings (low cut-off = 0.005; high cut-off = 0.500) to help delineate individual cells. Regions of interest (ROI) were selected by manually identifying cell-body-sized, high-contrast regions over the course of the timeseries, and contours were drawn to contain pixels from the ROI (*35*). Raw fluorescent values were exported for each ROI for the full timeseries. ROIs were further validated by inspecting temporal traces from individual ROIs for Ca^2+^ dynamics consistent with Ca^2+^ transients from individual cells, and only ROIs with clearly identified signals were included in further analyses (*18, 36–38*). We identified 586 ROIs from 6 mice. Custom MATLAB scripts were then used to align exported raw fluorescent values from imaging frames with scored sleep and wake states using annotations in the EEG & EMG data generated by the TTL pulses from Inscopix nVista imaging system.

To correct for slight decays in fluorescent signal across each recording, Ca^2+^ traces for each ROI were detrended by subtracting an exponential curve fit from each individual raw Ca^2+^ imaging trace using the ‘fit’ function from MATLAB’s Curve Fitting toolbox (*39*). The exponential curve is given by:

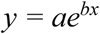

where *a* is the starting fluorescent value, *b* is the decay constant, *x* is time, and *y* is the fluorescent value at time *x*. The curve fit from all recordings across the timeseries were then averaged and added back to each trace to bring traces to a common baseline. Ca^2+^ values were then expressed as percent change from the median fluorescent value of the full timeseries:

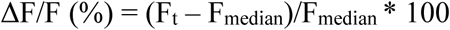

where F_t_ is the individual detrended fluorescent value at a specific timepoint and F_median_ is the median fluorescent value for the timeseries (*39, 40*). Event detection with a 0.5% prominence threshold was then performed using the ‘findpeaks’ function from MATLAB’s Signal Processing toolbox. Two conditions had to be met to identify an ROI’s frame as an event: 1) the value of that frame is larger than its two neighboring frames, and 2) the difference between that value and the value of the larger neighboring trough (prominence) is greater than 0.5% of the range of the entire trace. The amplitudes of these events were then sorted into deciles of the range. Typically, Ca^2+^ imaging studies set a threshold (e.g. 2 standard deviations) that an event or peak must cross to be counted as an event. However, this thresholding would eliminate the majority of astroglial Ca^2+^ events that occur during non-rapid eye movement sleep (NREMS) and rapid eye movement sleep (REMS) because these events are more modest compared to those in wake. Instead, all detected astroglial Ca^2+^ events were binned into deciles of the range by calculating the event’s amplitude as a percentage of the amplitude range of the trace across the full timeseries (i.e. across all states) rounded to the nearest 10%. To inspect Ca^2+^ dynamics during vigilance state transitions, Ca^2+^ data were extracted frame-by-frame for the 12 s before and after the transition. These transitions were removed from analysis for ΔF/F comparisons over time in Fig. 1G and fig. S3 as more state transitions were captured during ZT6 compared to other timepoints. The removal of these transitions allowed for comparisons of Ca^2+^ during steady state epochs. We did not include wake-to-REMS or REMS-to-NREMS transitions as these transitions rarely occur in mice. Pearson’s correlation coefficients were calculated from ΔF/F values across individual bouts of wake, NREMS, and REMS at ZT6. Vigilance state bouts used for the cross correlation analysis were 20 – 60 s long.

### Two-photon Imaging

#### Data acquisition

During experiments, mice were connected to a tether coupled to a nano connector. The tether was coupled to an interface box (S-Box, Tucker-Davis Technologies, Alachua, FL), transmitted EEG & EMG signals were amplified via a low impedance amp (PZ3-32, Tucker-Davis Technologies), and data were processed and recorded via the Z-Series 2 DSP High Performance Processor (RZ2-2, Tucker-Davis Technologies) and OpenEx software (Tucker-Davis Technologies). EEG and EMG data were digitized at 1017.3 Hz. EEG signals were high pass filtered at 0.3 Hz and low pass filtered at 30 Hz. EMG data were high pass filtered at 10 Hz and low pass filtered at 100 Hz. The system was also set up to receive and incorporate TTL pulses generated by the two-photon imaging system.

Prior to experiments, mice were habituated to increasingly prolonged periods of time in head restraint in the flat-floored, air-lifted Mobile HomeCage (Neurotar, Helsinki, Finland). During experiments, unanesthetized mice were head-fixed inside the Mobile HomeCage at least 30 min prior to imaging and then imaged using resonant scanning microscope UltimaIV running Prairie View acquisition software (Bruker Corporation, Billerica, MA). To excite, filter, collect, and detect GCaMP6f fluorescence changes in astrocyte somata and processes, we used an excitation wavelength of 920 nm (Chameleon II, Coherent Inc., Santa Clara, CA) with ET-GFP (FITC/CY2) emission filter cube (#49904, Chroma Technology Corporation, Bellows Falls, VT) with a 20x (1.0NA) XLUMPLFL20X W/IR-2 objective (Olympus, Shinjuku, Tokyo, Japan) and GaASP photo-multiplier tube (H7422-40, Hamamatsu Photonics KK, Hamamatsu City, Shizuoka, Japan). The imaged field of view was 277 x 277 µm, pixel dimensions were 0.188 x 0.188 µm, and rate of acquisition was 3.7 Hz. During frame acquisition, TTL pulses were sent from the two-photon microscope system to the Tucker-Davis Technology system and incorporated in the EEG & EMG data file for subsequent alignment of imaging frames and electrophysiological data.

#### Data analysis

The ImageJ plugin “Kalman Filter” was applied to collected image stacks to remove high gain noise and to recover faint image details. Imaged stacks were then visually inspected frame-by-frame which occasionally revealed motion artifacts along the x-and y-axes and required registration using the “Template Matching” ImageJ plugin. Image frames along the x- and y-axes that could not be registered or any that occurred along the z-axis were manually removed from analysis. These registered z-stacks could then be used to produce sum projection images (sums the slices in the stack) or maximum projection images (image displaying maximum pixel values from slices in the stack) in ImageJ. For selection criteria, first, astrocyte somata and processes were identified from high resolution, maximum projected image stacks. Second, individual GCaMP6f expressing astrocytes in which the somata and its processes were captured within the same image plane and did not overlap with other cells were identified for further analysis. For signal extraction, two separate ROIs were drawn per astrocyte, one circumscribing the cell soma and one defining the territory of its processes. With the territory of the astrocyte processes delineated, a sampling-grid containing 2.7 µm x 2.7 µm squares was overlaid. Data were extracted only from those ROI grid squares contained, in their entirety, within the circumscribed processes territory. From 4 mice, we identified n = 60 astroglial somata along with n = 6074 process ROIs.

Ca^2+^ imaging data were processed as described for the epifluorescent imaging data for each ROI and their respective process ROIs using the custom MATLAB scripts. To compare differences in the Ca^2+^ concentration of the somata and processes across vigilance states, we calculated frame-by-frame difference scores by subtracting the ΔF/F value of the soma from the ΔF/F value from each associated process ROI (ΔF/F_DIFF_ = ΔF/F_Process_ - ΔF/F_Soma_). While in head-restraint in the air-lifted cage, mice spent sustained periods of time in a “transitional state” characterized by high amplitude, high frequency EEG signals and low amplitude EMG signals. Ca^2+^ data collected during this transitional state were not included in analyses.

### Sleep Phenotyping

EEG and EMG data were collected with a Grass 7 polygraph system (Natus Medical Incorporated, Pleasanton, CA) via a light-weight cable for sleep phenotyping experiments in GSTIM1 WT and cKO mice. EEG & EMG signals were amplified and digitized at 256 Hz using VitalRecorder acquisition software (SleepSign for Animal, Kissei Comtec Co., LTD, Nagano, Japan). EEG and EMG data were high- and low-pass filtered at 0.3 and 100 Hz and 10 and 100 Hz, respectively.

All EEG and EMG data were scored using SleepSign for Animal. Wake, NREMS, and REMS were determined by visual inspection of the EEG waveform, EMG activity, and fast Fourier transform (FFT) analysis by an experimenter blinded to experimental conditions. Data were scored as wake, NREMS, or REMS with 4-s resolution as previously described (*2, 31*). Bout lengths were defined as ≥ 7 consecutive epochs (≥ 28 s) for wake and NREMS and ≥ 4 consecutive epochs (≥ 16 s) for REMS.

The EEG was subjected to FFT analysis to produce power spectra between 0 – 30 Hz with 0.5 Hz resolution. Delta (δ) was defined as 0.5 – 4 Hz and low delta as 0 – 1.5 Hz (*2*). For genotypic comparisons of EEG spectral data, each spectral bin was expressed as a percentage of the total power in baseline wake, NREMS, and REMS averaged across the 3 states. For hourly NREM delta power (i.e. NREM slow wave activity) analysis after SD, spectral values within the delta band or low delta band for each hour were normalized to the average NREM delta or low delta band value, respectively, from the last 4 h of the baseline light period (ZT8 – 11) and expressed as a percentage (*17*). EEG epochs containing visually-detected artifacts were excluded from spectral analyses. Two GSTIM1 cKO mice were removed from spectral analyses due to artifact.

### Statistical Analysis

Plots were generated in SigmaPlot (v11.0, Systat Software, Inc., San Jose, CA) and R (v3.6.0). For simplicity, outliers were not plotted with boxplots. Statistical analyses were performed using SPSS for Windows (IBM Corporation, Armonk, NY) unless otherwise noted. Data are presented as means ± standard error of the mean (s.e.m.). Normality of the data was determined with Shapiro-Wilk or Kolmogorov-Smirnov tests. Comparisons of baseline ΔF/F, amplitude deciles, events per epoch, Pearson correlation coefficients, and process-somata difference scores across vigilance states were made using Friedman tests. *Post-hoc* pairwise comparisons were made using Wilcoxon signed rank tests with Bonferroni corrections. The mice, mitml, and lme4 R packages were used to compare NREMS ΔF/F values and NREM EEG delta power across ZT0, 6, and 12 with multiple imputations using the 2l.pan mixed-effects model and 100 imputations. The imputed models were then fit using a linear mixed-effects model through the lmer function with individual mice as a random effect. *Post-hoc* pairwise comparisons of the fitted model were made with the emmeans function and Tukey correction. Comparisons of events/epoch and ΔF/F values between somata and processes were made with the Mann-Whitney U test. Comparisons of Ca^2+^ measures between baseline and SD days were made with Wilcoxon signed rank tests. For sleep phenotyping, a general linear model for repeated measures (RM) using time (hours or period) as the repeated measure and genotype (GSTIM1 WT vs. GSTIM1 cKO) as the between subjects factor was used when multiple measurements were made over time (i.e. time in stage, bout number, bout duration, hourly NREM delta power). Repeated measures were tested for sphericity, and a Greenhouse-Geisser correction was applied when appropriate. *Post-hoc* pairwise comparisons using Sidak corrections were performed when there were significant interaction effects or main effects of genotype. Baseline time in state and bout data RM comparisons were made over all time intervals during the full 24 h recording period (ZT0 - 23).

For SD experiments, RM comparisons were made over all time intervals during the full recovery period (ZT6 – 23) for time in state data. Comparisons of 2 h averages of NREM delta power post-SD were made with RM over ZT6 – 11 of the recovery phase. Comparisons of normalized EEG spectral power during baseline and after SD were made using RM comparison from 0 – 30 Hz with spectral power as the dependent variable and genotype as the between-subjects factor. If the RM comparisons yielded a significant result, unpaired Student’s t-tests with Benjamini-Hochberg correction were then used for comparisons of individual EEG spectral frequency bins with genotype as the grouping variable. Unpaired Student’s t-tests with genotype as the grouping variable were used to analyze total sleep time post-SD. An alpha level of 0.05 was used to indicate significance.

---

**Fig. S1.**
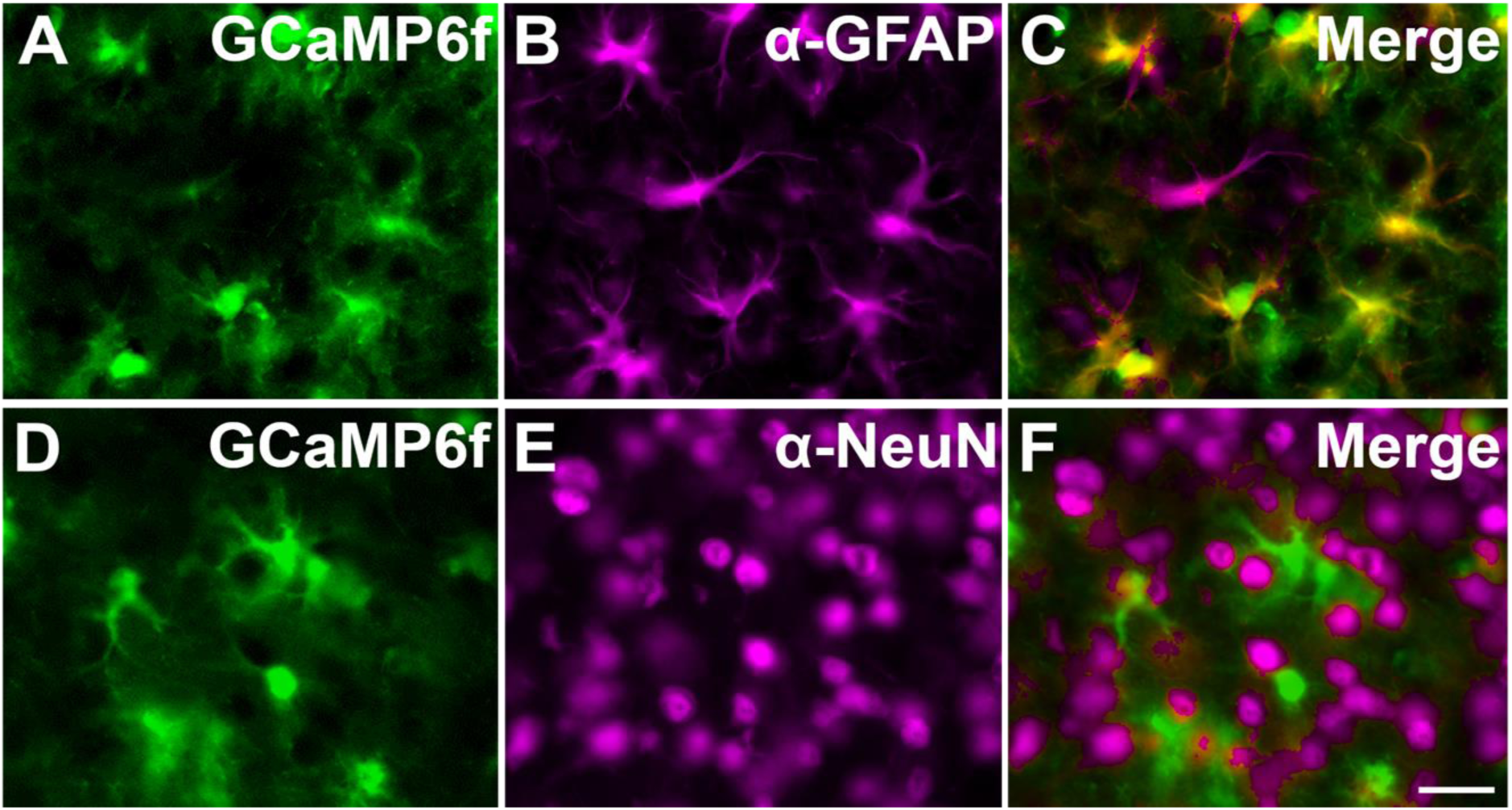
Selective GCaMP6f expression in astrocytes. Microinjection of AAV2/5 *GfaABC_1_D*-GCaMP6f in frontal cortex results in selective expression of GCaMP6f in GFAP^+^ astrocytes (**A** – **C**), but not in NeuN^+^ neurons (**D** – **F**). 40x objective; scale bar = 20 μm. GCaMP6f labeling is native (**A** and **D**) whereas astrocytes (GFAP; **B**) and neurons (NeuN, **E**) are immunolabeled.

**Fig. S2.**
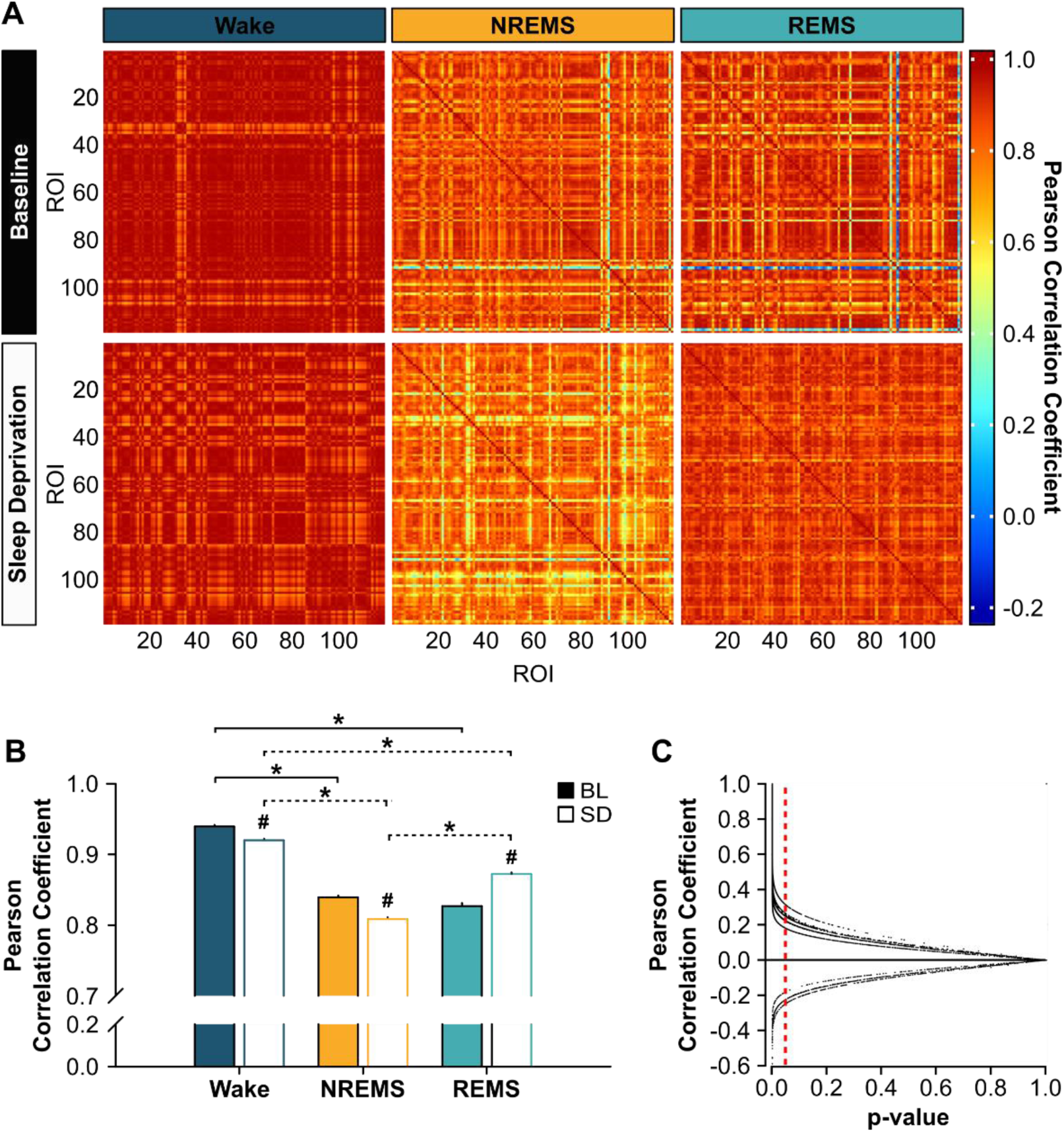
Pearson correlation coefficients of ΔF/F values during sleep and wake. (**A**) Pearson correlation coefficient matrices of ΔF/F values from the Inscopix microscope during wake (left), NREMS (middle), and REMS (right) under baseline (BL; top row) and sleep deprived (SD; bottom row) conditions from 119 regions of interest (ROI) from one representative mouse. Bout lengths shown in each matrix are as follows: BL, W = 20 s, NR = 60 s, R = 60 s; SD, W = 20 s, NR = 60 s, R = 60 s. (**B**) Mean Pearson correlation coefficients (n = 4 mice) during wake, NREMS, and REMS under BL (closed bars) and SD (open bars) conditions. Brackets denote comparisons between vigilance states for BL (solid) and for SD (dashed). # denotes comparisons between BL and SD within each vigilance state. (Between states: Friedman, BL, χ^2^ = 544.73, p < 0.001; SD, χ^2^ = 691.44, p < 0.001. BL vs. SD: Friedman, χ^2^ = 45.44, p < 0.001). p < 0.05. (**C**) Individual Pearson correlation coefficients from each mouse plotted against their respective p-value. Vertical, red, dashed line denotes a p-value of 0.05. Data points to the left of this line correspond to significant Pearson correlation coefficients.

**Fig. S3.**
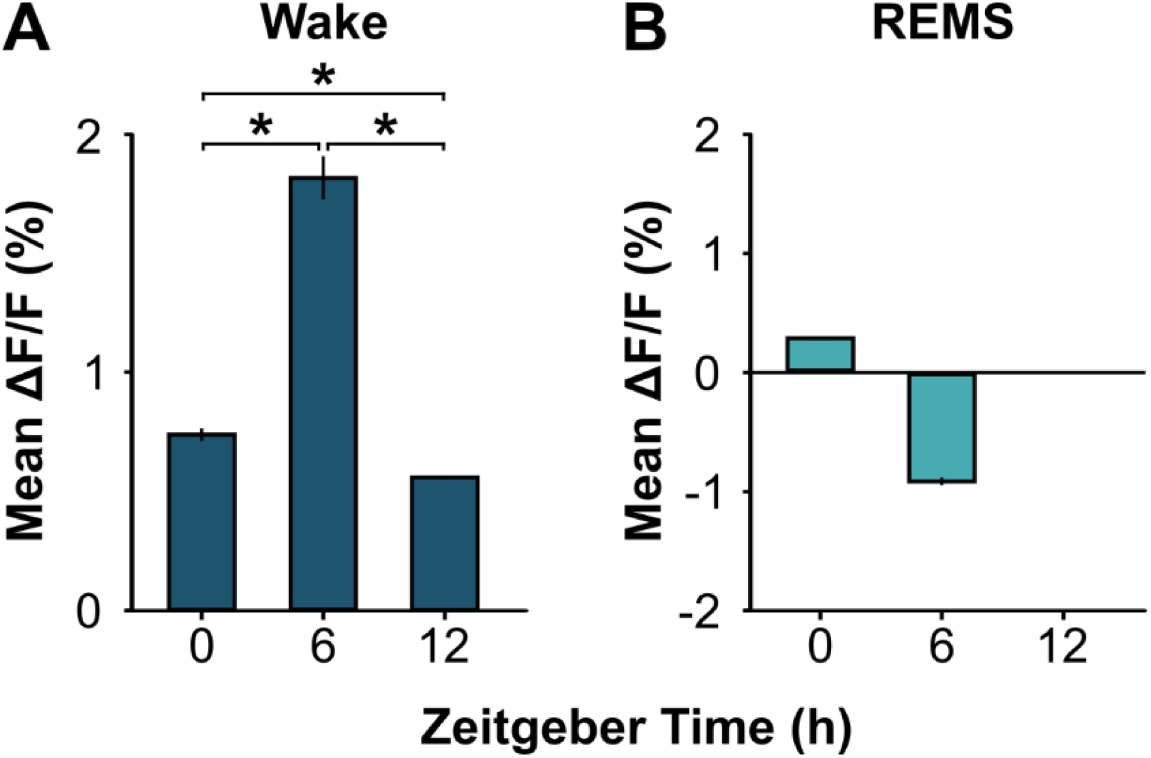
Mean ΔF/F during wake and REMS during ZT0, 6, and 12. Mean ΔF/F values from the Inscopix microscope during (**A**) wake (repeated measures ANOVA, time effect; F(1.133, 642.25) = 175.85, p < 0.001) and (**B**) REMS. REMS did not occur in recordings made at ZT12. Values are means ± s.e.m. *, different from ZT. p < 0.05.

**Fig. S4.**
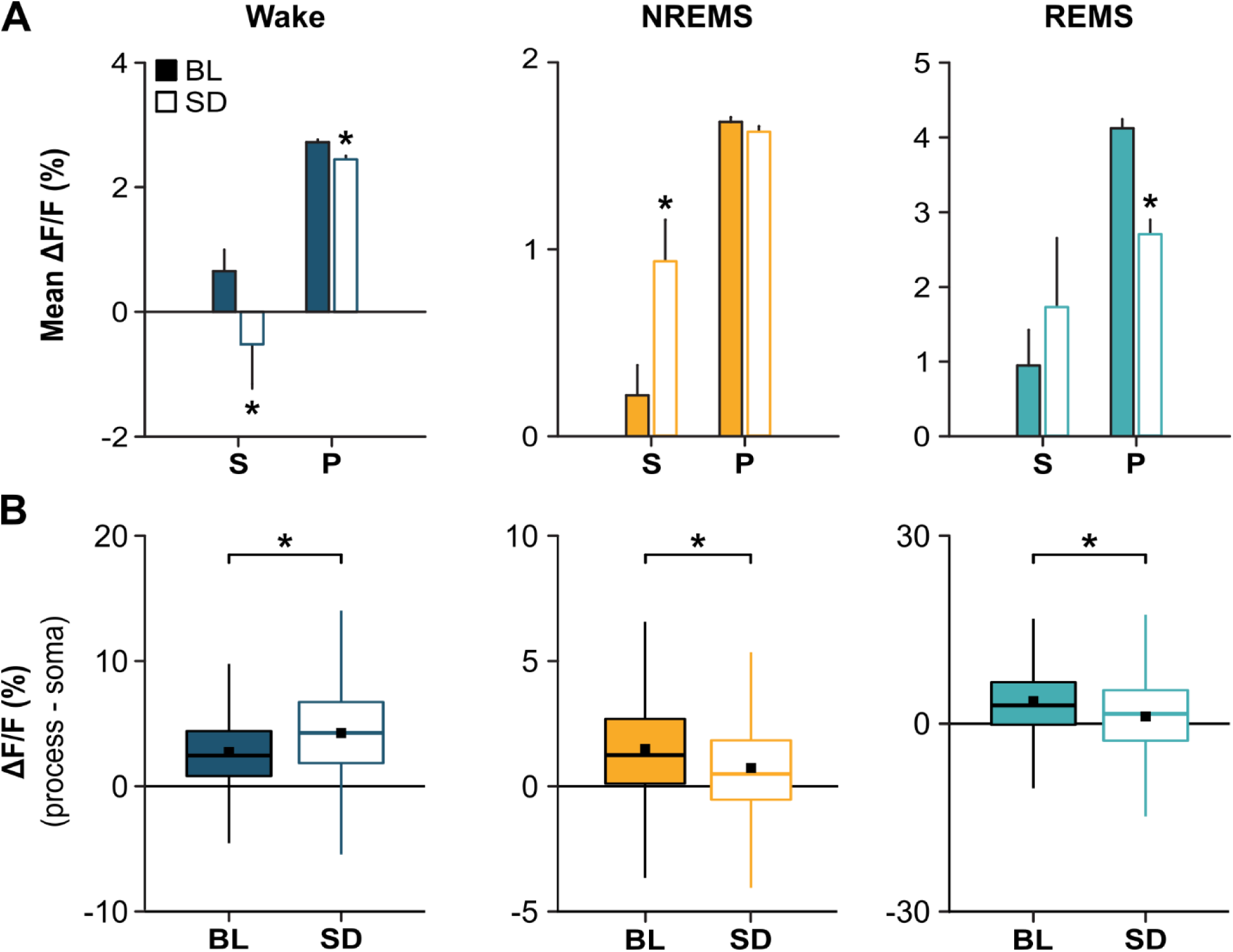
Astroglial Ca^2+^ dynamics in somata and processes at ZT12. (**A**) Mean ΔF/F values from the two-photon microscope at ZT12 during wake, NREMS, and REMS under baseline (BL; closed) and sleep deprived (SD; open) conditions for somata (S) and processes (P) (Somata: Wilcoxon signed rank, wake, Z = −3.39, p = 0.001; NREMS, Z = −3.15, p = 0.002; REMS, Z = −0.63, p = 0.532. Processes: Wilcoxon signed rank, wake, Z = −10.83, p < 0.001; NREMS, Z = −1.21, p = 0.224; REMS, Z = −8.53, p < 0.001). (**B**) Distribution of differences of ΔF/F values between processes and their soma. Black squares denote means (Wilcoxon signed rank, wake, Z = −24.45, p < 0.001; NREMS, Z = −27.06, p < 0.001; REMS, Z = −8.09, p < 0.001). *, different from BL. P < 0.05.

**Fig. S5.**
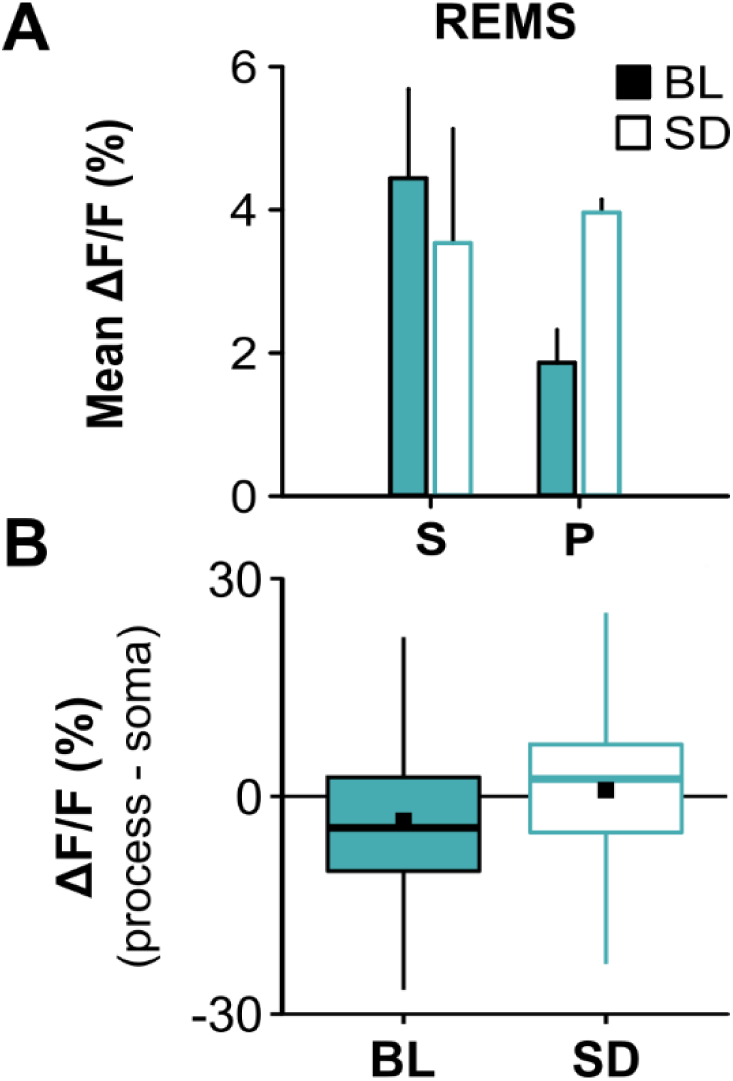
Astroglial Ca^2+^ dynamics in somata and processes in REMS at ZT6. (**A**) Mean ΔF/F values from the two-photon microscope at ZT6 during REMS under baseline (BL; closed) and sleep deprived (SD; open) conditions for somata (S) and processes (P). (**B**) Distribution of differences of ΔF/F values between the processes and somata. Black squares denote means.

**Fig. S6.**
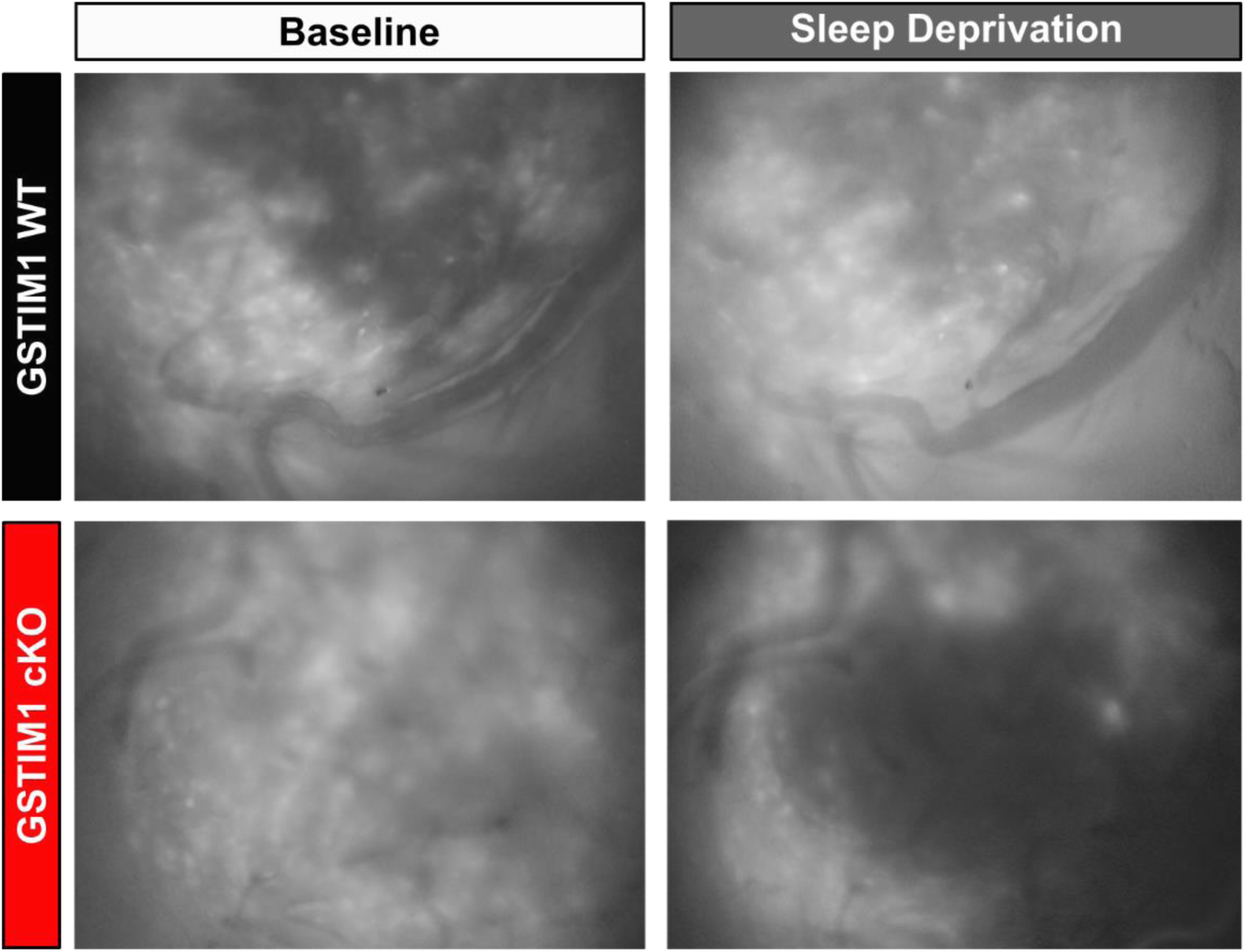
Astroglial Ca^2+^ is attenuated in GSTIM1 cKO mice. Representative maximum intensity z-stack projection images of GCaMP6f fluorescence from a GSTIM1 WT mouse (top) and a GSTIM1 cKO mouse (bottom) at ZT6 under baseline conditions (left) and after 6 h of sleep deprivation (right) from the Inscopix microscope.

**Fig. S7.**
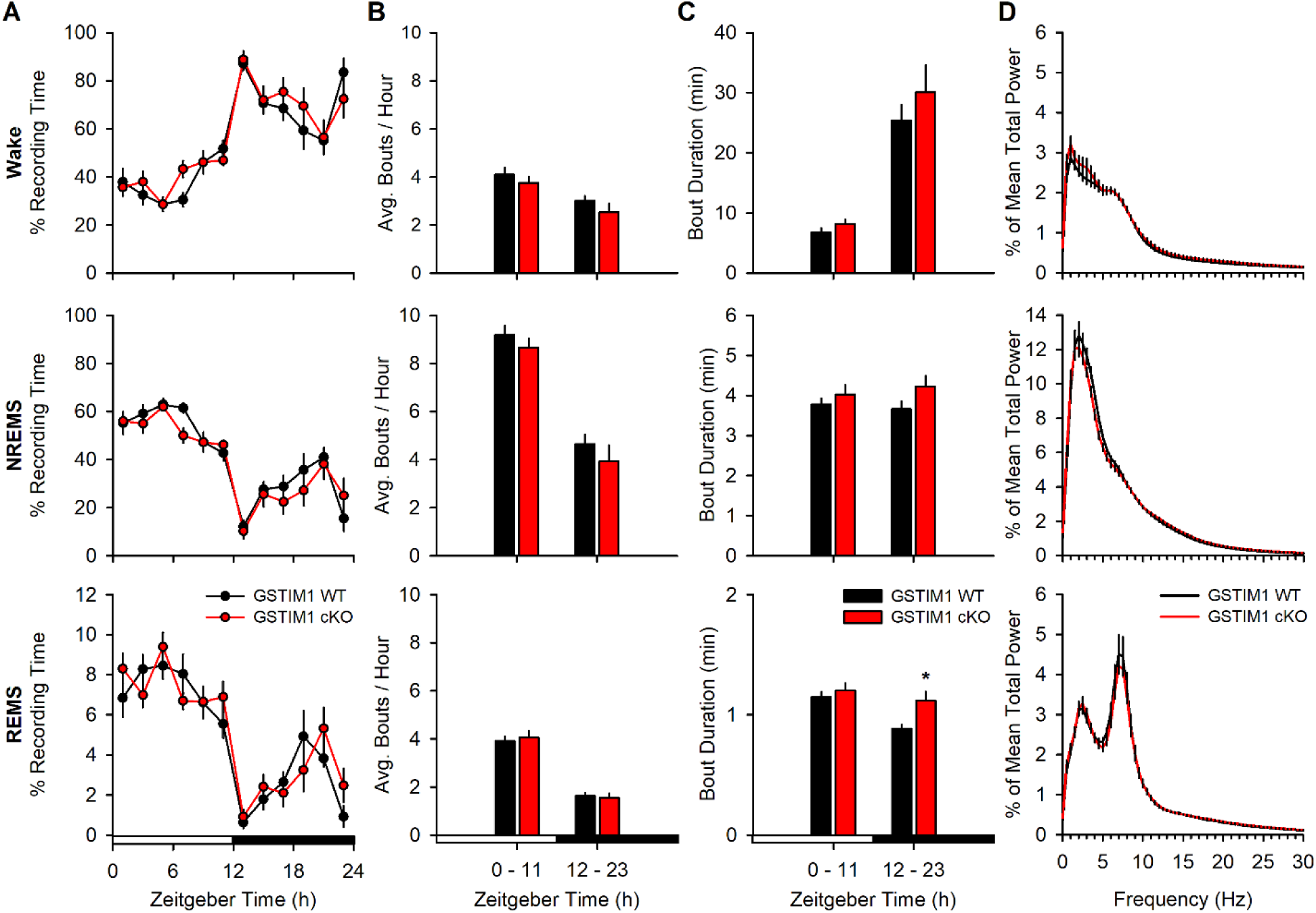
Baseline sleep-wake behavior is similar to wild type mice after conditional knock out (cKO) of STIM1 in astrocytes. Vigilance state data for wake (top), NREMS (middle), and REMS (bottom). (**A**) Time spent in wake, NREMS, and REMS expressed as a percentage of recording time in 2 h bins (repeated measures ANOVA). (**B**) Average number of state-specific bouts per hour shown in 12 h bins (repeated measures ANOVA). (**C**) Average bout duration (min) per hour shown in 12 h bins (repeated measures ANOVA, F(1, 15) = 7.70, p = 0.014). (**D**) 24 h state-specific EEG spectral power expressed as a percentage of the mean total power of all vigilance states (repeated measures ANOVA). Values are means ± s.e.m. *, different from GSTIM1 WT. p < 0.05.

**Fig. S8.**
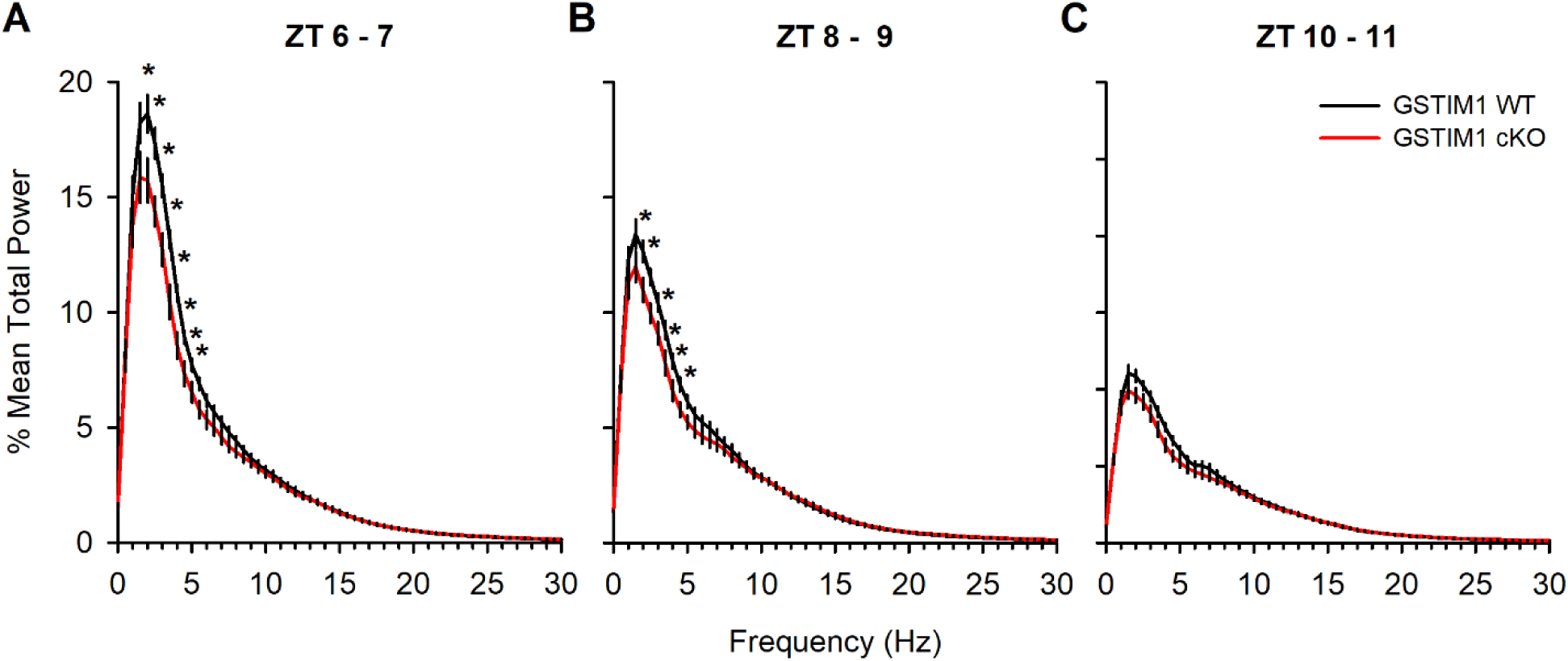
NREM EEG SWA is blunted in GSTIM1 cKO mice after 6 h sleep deprivation (SD). NREM EEG spectral power during the first 6 h of recovery sleep after 6 h SD shown in 2 h time blocks: (**A**) ZT 6 – 7 (repeated measures ANOVA, frequency x genotype, F(2.03, 26.41) = 4.72, p = 0.017), (**B**) ZT 8 – 9 (repeated measures ANOVA, frequency x genotype, F(2.03, 29.98) = 3.20, p = 0.049), and (**C**) ZT 10 – 11 (repeated measures ANOVA). Values are means ± s.e.m. expressed as a percentage of the mean total baseline power of all states. *, genotypic differences. p < 0.05.

**Movie S1.**

**Astroglial Ca^2+^ dynamics during wake, NREMS, and REMS.** Astroglial Ca^2+^ imaging captured with the head-mounted, epifluorescent microscope (Inscopix) in a freely behaving mouse. Movie is a ΔF/F video that corresponds to traces shown in Fig. 1B sped up 8x. Fluorescent values are within the detectable range.

**Movie S2.**

**Astroglial Ca^2+^ dynamics in somata and process during sleep and wake.** Astroglial Ca^2+^ imaging captured with two-photon microscopy in an unanesthetized, head-restrained mouse. Movie corresponds to traces shown in Fig. 2B sped up 8x.

